# COI metabarcoding primer choice affects richness and recovery of indicator taxa in freshwater systems

**DOI:** 10.1101/572628

**Authors:** Mehrdad Hajibabaei, Teresita M. Porter, Michael Wright, Josip Rudar

**Affiliations:** Centre for Biodiversity Genomics @ Biodiversity Institute of Ontario & Department of Integrative Biology, University of Guelph, Guelph, Ontario, Canada, N1G 2W1; Natural Resources Canada, Canadian Forest Service, Great Lakes Forestry Centre, 1219 Queen St. East, Sault Ste. Marie, Ontario, Canada, P6A 2E5

**Keywords:** metabarcoding, biomonitoring, cytochrome c oxidase (COI), primers, indicator species, Ephemeroptera, Plecoptera, Trichoptera, Chironomidae

## Abstract

DNA-based biodiversity analysis has gained major attention due to the use of high throughput sequencing technology in approaches such as mixed community or environmental DNA metabarcoding. Many cytochrome c oxidase subunit I (COI) primer sets are now available for such work. The purpose of this study is to look at how COI primer choice affects the recovery of arthropod richness, beta diversity, and recovery of site indicator taxa in benthos kick-net samples typically used in freshwater biomonitoring. We examine 6 commonly used COI primer sets, on samples collected from 6 freshwater sites. Richness is sensitive to primer choice and the combined use of additional multiple COI amplicons recovers higher richness. Thus, to recover maximum richness, multiple primer sets should be used with COI metabarcoding. Samples consistently cluster by site regardless of amplicon choice or PCR replicate. Thus, for broadscale community analyses, overall beta diversity patterns are robust to COI marker choice. Additionally, the recovery of traditional freshwater bioindicator assemblages such as Ephemeroptera, Trichoptera, Plectoptera, and Diptera may not fully capture the diversity of broadscale arthropod site indicators that can be recovered from COI metabarcoding. Based on these results, studies that use different COI amplicons may not be directly comparable. This work will help future biodiversity and biomonitoring studies develop not just standardized, but optimized workflows that maximize taxon-detection or order taxa along gradients.

## Introduction

DNA-based biodiversity analysis has gained major attention due to the use of high throughput sequencing technology in approaches such as mixed community or environmental DNA metabarcoding (Hajibabaei, Shokralla, Zhou, Singer, & Baird, 2011; Taberlet, Coissac, Pompanon, Brochmann, & Willerslev, 2012). Data generation typically involves DNA extraction from an environmental sample such as water, soil or collected biomass (e.g. benthic kicknet, malaise trap) followed by PCR amplification of one or more taxonomic markers such as the COI DNA barcode region and subsequent high throughput sequencing and bioinformatic analysis of marker gene sequences. Resulting sequences are then assigned to sequence clusters (Operation Taxonomic units, OTU; Exact Sequence Variants, ESV) and/or taxonomic names (Callahan, McMurdie, & Holmes, 2017). Sequence clusters and taxonomic lists obtained are used in various statistical analyses for assessing different aspects of biodiversity such as species richness or distribution, community composition, and functional diversity(Porter & Hajibabaei, 2018c). In practice, these questions are often geared towards identifying assemblages or specific target taxa. Biodiversity information gained can contribute to ecological investigations and applications such as biomonitoring as part of environmental assessment programs (Baird & Hajibabaei, 2012; Leese et al., 2018).

A major step in obtaining sequence data from environmental samples involves PCR amplification of target marker gene(s). An important consideration in this multi-template PCR step is the choice of primer sets. It has been shown that primers can differentially bind to template DNA and can result in both qualitative and quantitative biases (Suzuki & Giovannoni, 1996; Polz & Cavanaugh, 1998; L. J. Clarke, Soubrier, Weyrich, & Cooper, 2014). Most previous metabarcoding studies use a single primer set for generating sequence data from communities, but there is ample evidence that multiple amplifications with different primer sets can provide better biodiversity coverage from environmental samples (ex. J. Gibson et al., 2014).

Current methods for biomonitoring especially in freshwater typically rely on key taxonomic groups that are considered ecological bioindicators. For example, Ephemeroptera, Plecoptera, Trichoptera are known to be sensitive to water pollution whereas Chironomidae (Diptera) have been shown to be tolerant to high levels of pollution (Buss, Baptista, Silveira, Nessimian, & Dorvillé, 2002; Bonada, Prat, Resh, & Statzner, 2006). Here we collectively refer to this assemblage as the EPTC. Because of the difficulties associated with morphological identification of larval samples from benthos, samples are generally identified to family or genus level. Sorting and identifying individual samples from benthos poses a serious challenge in executing large-scale biomonitoring programs. With the advancement of genomics methods such as DNA metabarcoding, sequence data is generated from whole communities without the need to isolate individuals. Therefore, the analysis can go beyond the target assemblages such as EPTC.

The objective of this study was to test the performance of several newly published COI metabarcode primers to detect freshwater benthic invertebrates. We wanted to determine the impact of primer choice on several components of diversity: richness, beta diversity, and recovery of bioindicators. We tested a total of 6 partial COI metabarcode amplicons, including the two amplicons we have used routinely for freshwater invertebrate monitoring.

## Methods

### Field methods

Six benthic invertebrate communities were sampled from shallow streams across the City of Waterloo (Ontario, Canada) using a modified travelling kick-and-sweep technique outlined in the Ontario Benthos Biomonitoring Network protocol (Jones, Somers, Craig, & Reynoldson, 2007) (Table S1). Briefly, wetted width was measured and used to calculate the number of return trips required to sample a 10m transect of the stream specifically targeting a riffle habitat. Prior to sampling D-nets were decontaminated by soaking them in a 10% bleach solution for 15 min, rinsing with tapwater, and drying them overnight, A clean 500 µm mesh D-net was held downstream to the person sampling, with the opening of the net facing the person sampling. Substrate was disturbed by kicking the substrate at a constant effort for 3 minutes across the 10 m transect dislodging invertebrates and allowing the flowing water to guide the dislodged macroinvertebrates into the net. The samples were transferred from the net to a clean 1 L polyethylene bottle, preserved with 80% ethanol and stored at - 20°C until further processing in the lab.

### Molecular Biology Methods

#### DNA Extraction

Samples were homogenized separately in a clean blender (decontaminated thoroughly with Eliminase solution (Decon Labs: King of Prussia, PA, USA) (Black and Decker Model: BL2010BGC), distributing 50 mL of the homogenate into 6 sterile conical tubes for each sample. Samples were centrifuged at 2400 x g for 2min to collect homogenate at the bottom of the tube, and excess preservative ethanol was removed. Samples were covered and incubated at 65°C until residual ethanol was evaporated (roughly 5-8 hours). DNA was extracted using Qiagen’s DNeasy PowerSoil kit (Toronto, Canada. Product Ref: 12888). Samples were lysed overnight (∼ 15 hr). Following lysis, samples were extracted according to the manufacturer’s protocol, eluting with 30 μL molecular biology grade water. All extractions included a negative control where no sample was included.

#### Polymerase Chain Reaction

The six amplicons from CO1 barcode region used in this study are shown in Figure 1. The primers were aligned against the *Drosophila yakuba* COI barcode region obtained from GenBank accession X03240 using Mesquite v3.10 (Maddison & Maddison, 2015). COI secondary structure, 6 alpha helices, from *Bos taurus* were obtained from UniProt accession P00396. All samples were amplified for six primer sets according to their published amplification regime (Table 2) with the exception that a two-step PCR was used for all reactions (first PCR using untailed primers, second PCR using Illumina adapter-tailed primers), even if a one-step PCR was used in the original protocol. PCRs were run in duplicate with a negative control. Amplification success was confirmed through gel electrophoresis (not pictured).

**Table 1.**
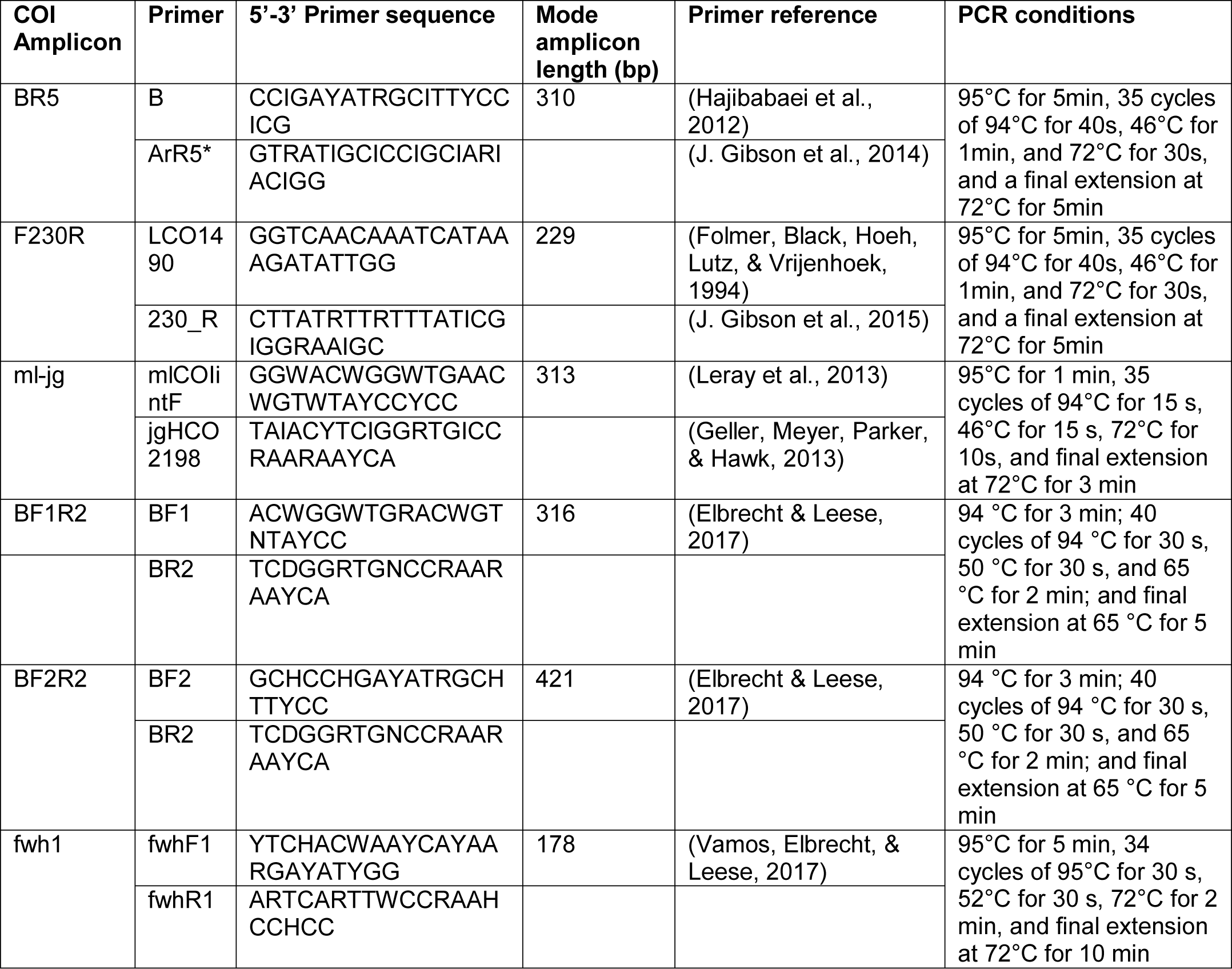
COI amplicons used in this study.

**Table 2.**
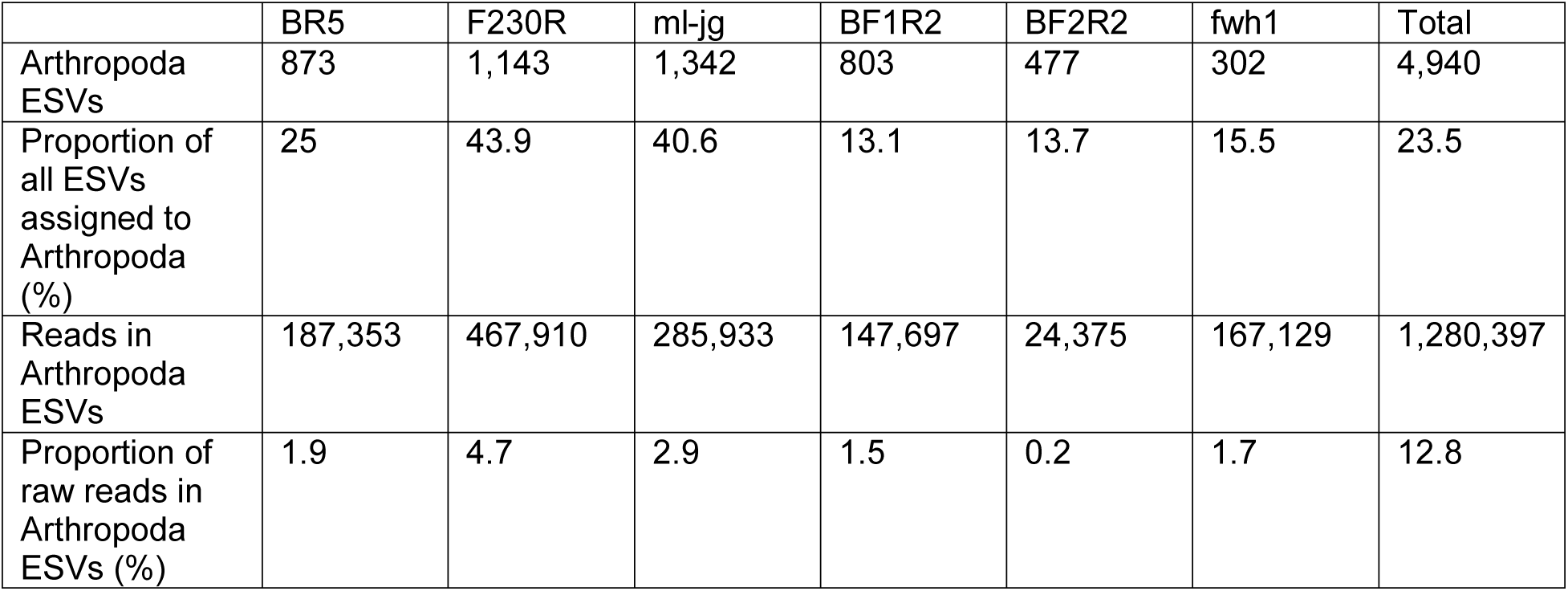
Arthropoda ESV and read counts vary by COI amplicon.

|  | BR5 | F230R | ml-jg | BF1R2 | BF2R2 | fwh1 | Total |
| --- | --- | --- | --- | --- | --- | --- | --- |
| Arthropoda<br>ESVs | 873 | 1,143 | 1,342 | 803 | 477 | 302 | 4,940 |
| Proportion of<br>all ESVs<br>assigned to<br>Arthropoda<br>(%) | 25 | 43.9 | 40.6 | 13.1 | 13.7 | 15.5 | 23.5 |
| Reads in<br>Arthropoda<br>ESVs | 187,353 | 467,910 | 285,933 | 147,697 | 24,375 | 167,129 | 1,280,397 |
| Proportion of<br>raw reads in<br>Arthropoda<br>ESVs (%) | 1.9 | 4.7 | 2.9 | 1.5 | 0.2 | 1.7 | 12.8 |

**Figure 1.**
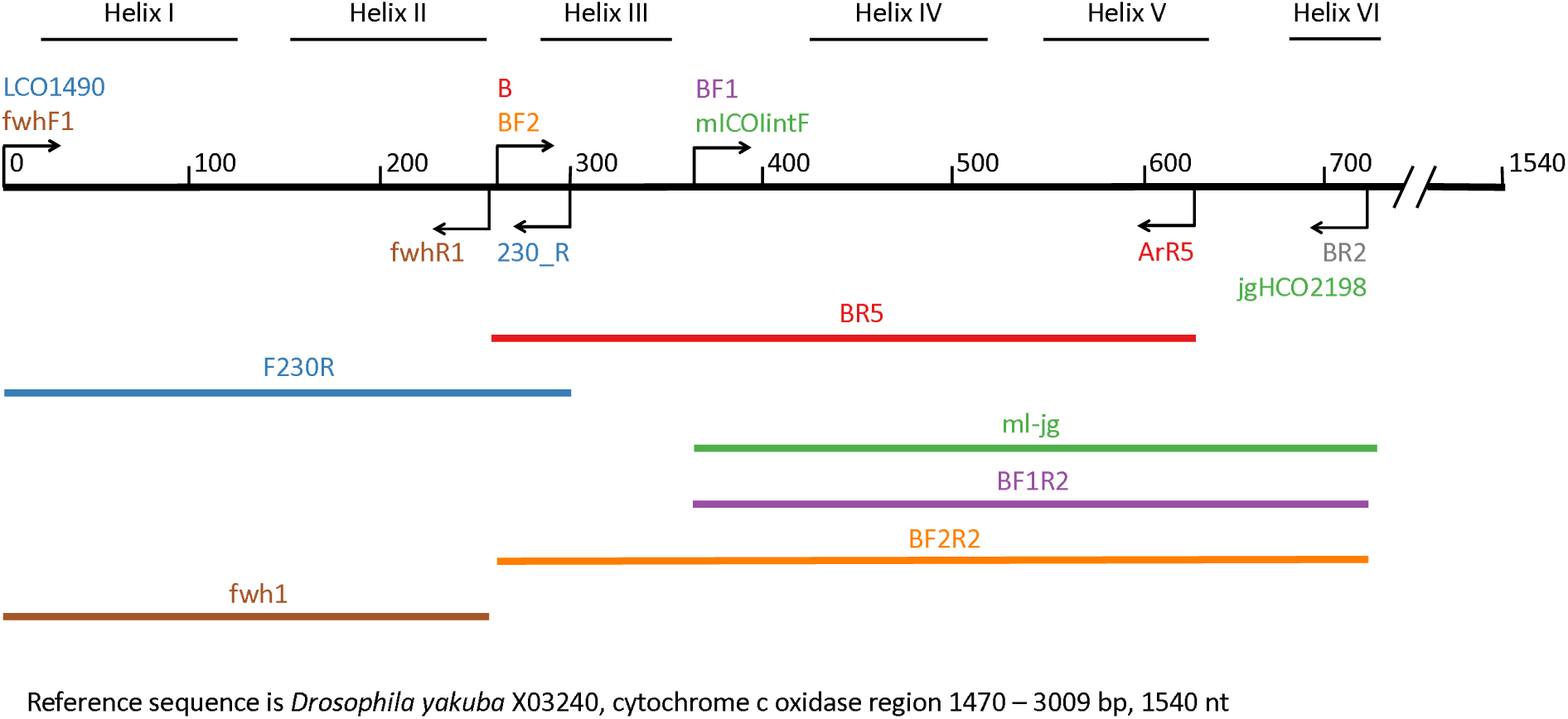
Map of primers and amplicons tested in this study. The reference sequence shown in black is *Drosophila yakuba*, cytochrome c oxidase region 1470-3009 bp (1540 nt). Secondary structure is shown for reference, comprised of 6 alpha helices in the standard DNA barcode region shown here.

Amplicons were purified using a MinElute PCR Purification kit, quantified on a TBS-380 Mini-Fluorometer (Turner Biosystems Sunnyvale California, United States) using a Quant-iT PicoGreen dsDNA assay (Invitrogen Waltham Massachusetts, United States Product Ref: P11496). The concentration of each purified sample was normalized across samples and primer sets were pooled for each sample. Although all primers were tested using the PowerSoil kit, samples extracted with the NucleoSpin Tissue Kit and amplified for BR5 and F230R primer sets were also sequenced as a comparison for their past use (J. Gibson et al., 2015). PCR replicates were sequenced separately from each other. Pooled samples were indexed using Illumina’s Nextera index kit (San Diego, California, United States Product Ref: FC-121-1011). All indexed samples were pooled, purified through magnetic bead purification, quantified using the PicoGreen dsDNA assay, and average fragment length for the library was determined on an Agilent Bioanalyzer 2100 (Santa Clara, California, United States. Product ref:G2939BA) using the Agilent DNA 7500 assay chip (Product Ref: 5067-4627). The library was diluted then sequenced using Illumina’s MiSeq v3 sequencing chemistry kit (2×300 cycle. Product Ref: MS-102-3003) on an Illumina MiSeq, comprising approximately half of a sequencing run.

### Bioinformatic processing

Raw sequences were processed with the SCVUC COI metabarcode pipeline v2.1 available from Github at https://github.com/EcoBiomics-Zoobiome/SCVUC_COI_metabarcode_pipeline. The acronym SCVUC stands for the major programs or algorithms used for bioinformatic processing: “S” – SEQPREP, “C” – CUTADAPT, “V” – VSEARCH, “U” – UNOISE, “C” – COI Classifier. Briefly, this semi-automated pipeline is described below. Jobs were spread across multiple cores using GNU Parallel (Tange, 2011). Raw compressed fastq Illumina read files were paired using SeqPrep specifying a minimum Phred score of 20 at the ends of the reads and an overlap of at least 25 bp (St. John, 2016). The following steps were conducted separately for each of the 6 amplicons tested in this study. Primers were trimmed using CUTADAPT v1.14 and reads were retained if they were at least 150 bp long after trimming, had a minimum Phred score of 20 at the ends of the reads, and contained no more than 3 N’s. CUTADAPT was also used to convert fastq files to FASTA files (Martin, 2011). The individual sample files were combined into a single file for global ESV generation. VSEARCH v2.4.2 was used to dereplicate the data (get the unique reads) using the –derep_fulllength option (Rognes, Flouri, Nichols, Quince, & Mahé, 2016). The USEARCH v10.0.240 unoise3 algorithm was used to denoise the reads (Edgar, 2016). This involved the removal of any contaminating PhiX reads (carry over from Illumina sequencing), prediction and removal of sequences with errors, removal of putative chimeric sequences, and removal of rare sequences. We defined rare sequences to be those clusters comprised of less than 3 reads (singletons and doubletons) (Brown et al., 2015; Tedersoo et al., 2010). We used this set of exact sequence variants (ESVs) as a reference, and all primer trimmed reads were mapped to this reference set with an identity of 1.0 (100% sequence similarity) to generate a sample x ESV table. The COI Classifier v3.2, that uses the Ribosomal Database Project naïve Bayesian classifier v2.12 with a custom COI reference set, was used to taxonomically assign the ESVs (Porter & Hajibabaei, 2018a; Wang, Garrity, Tiedje, & Cole, 2007). Taxonomic assignments were mapped to ESVs detected in each sample with a custom Perl script. The final taxonomy table for each primer was concatenated.

### Data analysis

The final taxonomy table above was formatted in R v3.4.3 in RStudio v1.1.419 (RStudio Team, 2016; R Core Team, 2017). Custom scripts are available from GitHub at URL. Data was summarized multiple taxonomic ranks. High confidence taxonomic assignments were retained by filtering for bootstrap support cutoffs >= 0.30 at the genus rank and >= 0.20 at the family rank. Using these cutoffs ensures that 95-99% of the taxonomic assignments are correct, assuming our query taxa are in the reference database (Porter & Hajibabaei, 2018a). We retained taxa at the species rank with a bootstrap support cutoff >= 0.70. Assuming our query species are present in the reference database, this should ensure that at least 95% of species level assignments are correct. To check whether we had sufficient sequencing depth, we used the package VEGAN v2.5-2 to plot rarefaction curves using the ‘rarecurve’ function (Oksanen et al., 2018). Curves that reach a plateau show saturated sequencing. To account for variable library sizes, reads/library were rarefied down to the 15^th^ percentile library size using the ‘rrarefy’ function (S. Weiss et al., 2017).

We compared average richness recovered from each amplicon using the VEGAN ‘specnum’ function and total richness was plotted with ggplot2 (Wickham, 2009). Richness data was checked for normality using visual distribution plots (ggdensity and ggqqplot, not shown) as well as using the Shapiro-Wilk test of normality (W=0.97, p=0.36) and this data was treated as normally distributed in comparisons (Shapiro & Wilk, 1965). We compared average richness using paired t-tests with the Holm adjustment for multiple comparisons.

There is uncertainty in how to interpret read abundance from arthropod metabarcoding studies due to unexpected template to product ratios after PCR due to stochasticity and GC content (Polz & Cavanaugh, 1998) as well as the effect of primer bias and body size variation across life stages and species that can vary by orders of magnitude and affect recovery (Elbrecht & Leese, 2015). As a result, we chose to transform read abundance into presence-absence data for all subsequent analyses. We checked for correlations in the presence-absence of ESVs recovered from DNA extractions processed with soil or tissue kits as well as between two PCR replicates using the ‘cor’ and ‘corrplot’ functions in R (Wei & Simko, 2017).

Indicator species can be used as a proxy to indicate differences among sites or conditions (De Cáceres & Legendre, 2009). For example, in freshwater systems, the diversity of EPTC taxa have been used as water quality indicators (Emilson et al., 2017). To test whether the recovery of indicator taxa depends on the COI amplicon used for the analysis, we performed indicator species analyses using the INDICSPECIES package in R and the ‘multipatt’ function with default settings (De Cáceres & Legendre, 2009). The six sites were used as groups for the analysis at multiple ranks and significant site indicators were selected if the resulting p-value was <= 0.05. We tested how often traditional EPTC are recovered with COI metabarcode data by repeating the analysis using all arthropod ESVs and just the ESVs assigned to EPTC.

To test whether sample clusters are affected by COI amplicon choice or PCR replicate, we used non-metric multidimensional scaling. Plots were created using the vegan ‘metaMDS’ function using the default settings with two dimensions (scree plot not shown) and dissimilarities were calculated using the method ‘bray’ for binary data (Sorensen dissimilarity) and plotted with ggplot. Goodness of fit was calculated using the VEGAN ‘goodness’ function. To check whether we had homogenous dispersion of dissimilarities, an assumption of permutational multivariate analysis of variance (PERMANOVA), we created a dissimilarity matrix with the VEGAN ‘vegdist’ function, then calculated beta dispersion using the ‘betadisper’ function in R. We tested for significant heterogeneity using analysis of variance (ANOVA) in R. We checked for significant interactions among sites, amplicons, and replicates as well as the significance of group clusters with PERMANOVA using the VEGAN ‘adonis’ function with 999 permutations.

## Results

A total of 9,980,584 x 2 paired-end reads were generated for this study and they have been deposited to the NCBI SRA: accession numbers # (Table S2). After pairing and primer trimming we retained a total of 7,619,108 reads. A summary of ESV counts for all taxa are shown in Table S3. About 23% of raw reads were retained in the denoised set of ESVs whereas the difference was removed during denoising as putative sequence errors, chimeras, PhiX contamination, or rare singletons and doubletons. About 24% of all ESVs were taxonomically assigned to Arthropoda taxa and the final Arthropoda ESV counts are shown in Table 2. When 6 COI primer pairs are compared, F230R ESVs contained the highest proportion of Arthropoda ESVs (43.9%) and contained the highest proportion of raw reads (4.7%). About 13% of raw reads were retained in this final set of Arthropoda ESVs. Out of all the Arthropoda taxonomic assignments, 11% of unique species, 15% of genera, and 26% of families were considered high confidence assignments (Table S4). The proportion of raw reads represented in these high confidence Arthropoda assignments was 7% for species, 8% for genera, and 10% for families.

Rarefaction curves show that at each rank, all samples reached a plateau, indicating that we had sufficient sequencing coverage for these samples (Figure S1). The proportion of taxa that are arthropods and the proportion of arthropods that are EPTC are shown in Figure S2. The average Arthropoda richness was not significantly different across the pairwise amplicon comparisons (pairwise t test, p > 0.05) and there was substantial variation in richness across sites (Figure S3). The total number of unique Arthropoda taxa were compared across COI amplicons (Figure S4) and the amplicon that detects the most unique taxa varied depending on the taxonomic resolution of the results. At the ESV rank, the ml-jg amplicon recovered the highest richness. We also note that the presence-absence of ESVs are positively correlated whether tissue or soil DNA kits are used for extraction (Figure S5) and across 2 PCR replicates (Figure S6).

To test the effect of using multiple COI amplicons on richness, we pooled increasing numbers of combined amplicons. We show that using a multi-amplicon approach can detect greater richness than using any single amplicon alone (Figure 2). In this study, ESV richness increases linearly as amplicons are added whereas species richness reaches a plateau when at least 4 amplicons are combined. In some cases, multiple combinations of amplicons recover equivalent richness. Due to limitations in the underlying reference sequence database, it is likely that species richness will also increase as additional reference taxa are added so that more ESVs can be assigned with high-confidence (Porter & Hajibabaei, 2018b).

**Figure 2.**
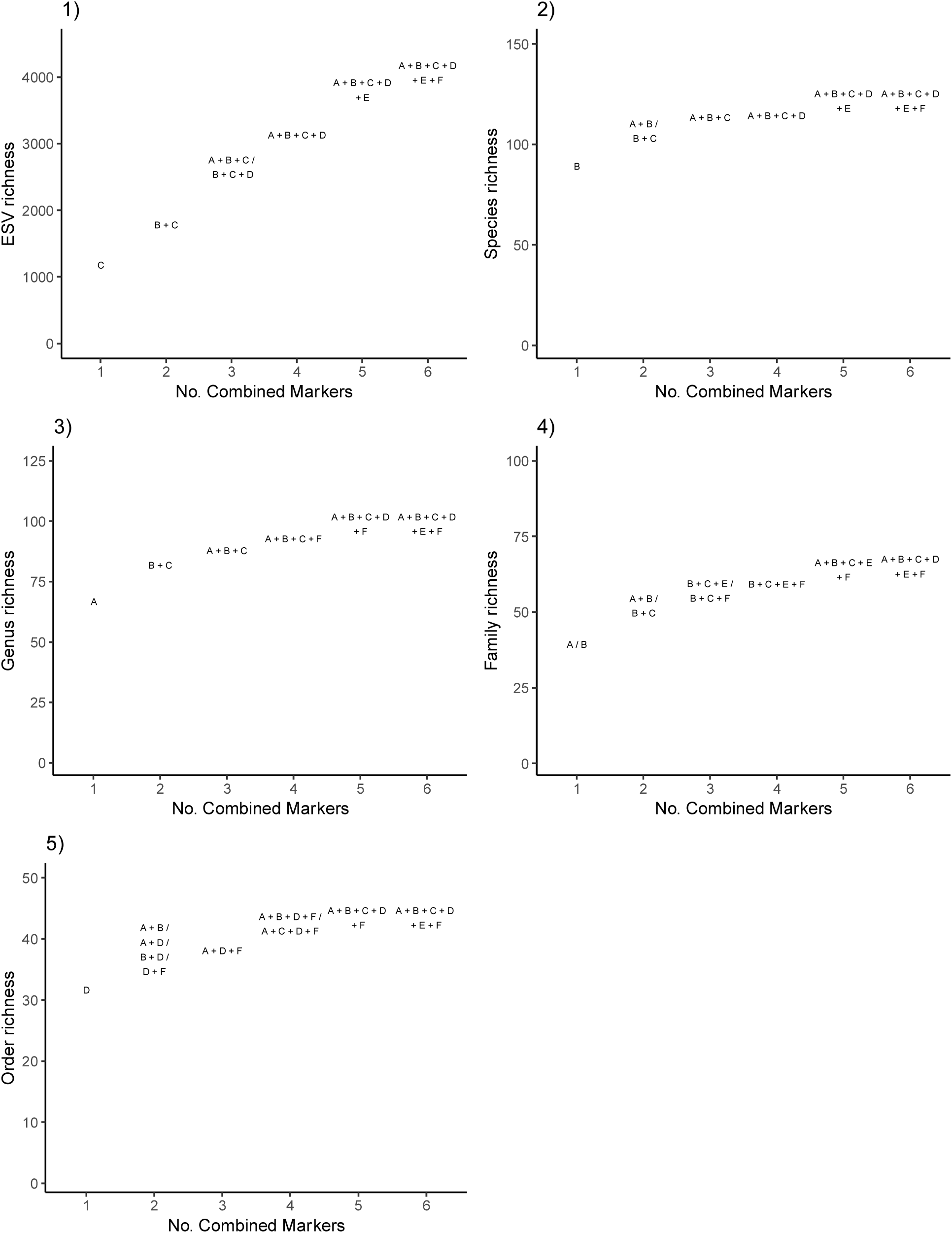
ESV richness continues to increase as COI amplicons are added but species - order richness reaches a plateau. For the primer comparison experiment that used the soil DNA extraction kit, we pooled the results from the 6 sites and show the top COI amplicon combinations that detected the greatest richness. We report the recovered richness when up to 6 amplicons are combined at the 1) ESV, 2) species, 3) genus, 4) family, and 5) order ranks. ESV = exact sequence variant; A = BR5; B = F230R; C = ml-jg; D = BF1R2; E = BF2R2; F = fwh1.

We also looked at how the recovery of broad-based indicator taxa from across the Arthropoda and more traditional freshwater indicator taxa from the EPTC varied with amplicon choice (Figure 3). Overall, more site indicator taxa were recovered from across the Arthropoda compared with limiting analyses to the EPTC. Generally, the amplicon that recovers the greatest number of site indicators varies according to the taxonomic resolution of the analysis. At the ESV rank, BF1R2 recovers the greatest number of broadscale site indicator taxa and F230R specifically recovers the greatest number of EPTC indicator taxa. Since the subset of indicator taxa presented for the species to family ranks only represents the portion of the ESVs assigned with high confidence, rank specific results may change over time as reference databases better represent local taxa (Porter & Hajibabaei, 2018b). We further explored the identities of non-traditional freshwater indicator taxa by looking at the taxonomic distribution of the broadscale indicator species and how this varied for each amplicon (Figure 4). Indicator taxa from the Elmidae (riffle beetles), Limoniidae (crane flies), and Simuliidae (black flies) were detected in addition to the expected indicator species from the Ephemeroptera and Trichoptera.

**Figure 3.**
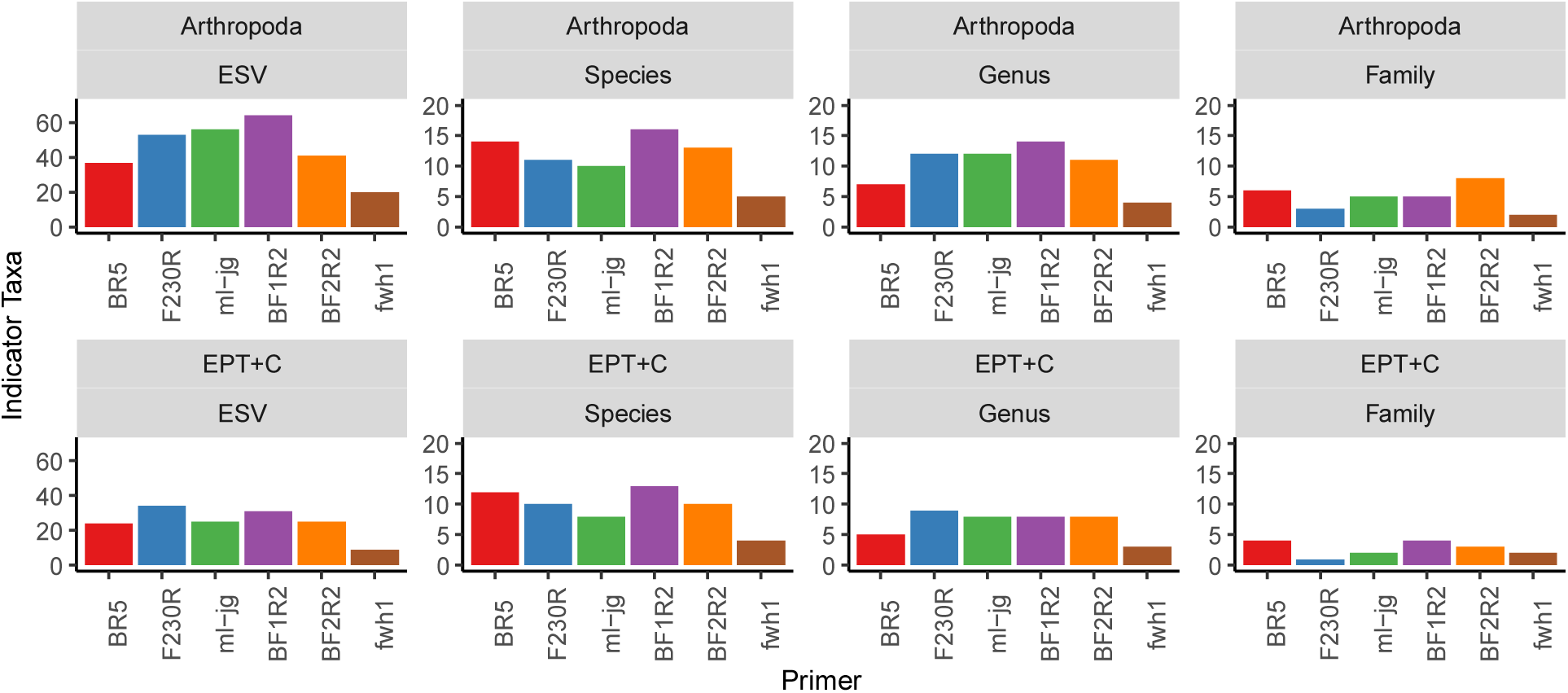
Ephemeroptera, Plecoptera, Trichoptera, and Chironomidae comprise a subset of total number of site indicator taxa drawn from across the Arthropoda. An indicator taxon analysis was used to determine the number of taxa that could distinguish among 6 sampled sites. In the top panel, the number of broadscale indicator taxa from across the Arthropoda are shown. In the bottom panel, the number of typical freshwater indicator taxa from the EPT+C are shown. This analysis was based on normalized data. ESV = exact sequence variant; EPTC = Ephemeroptera, Plecoptera, Trichoptera, Chironomidae.

**Figure 4.**
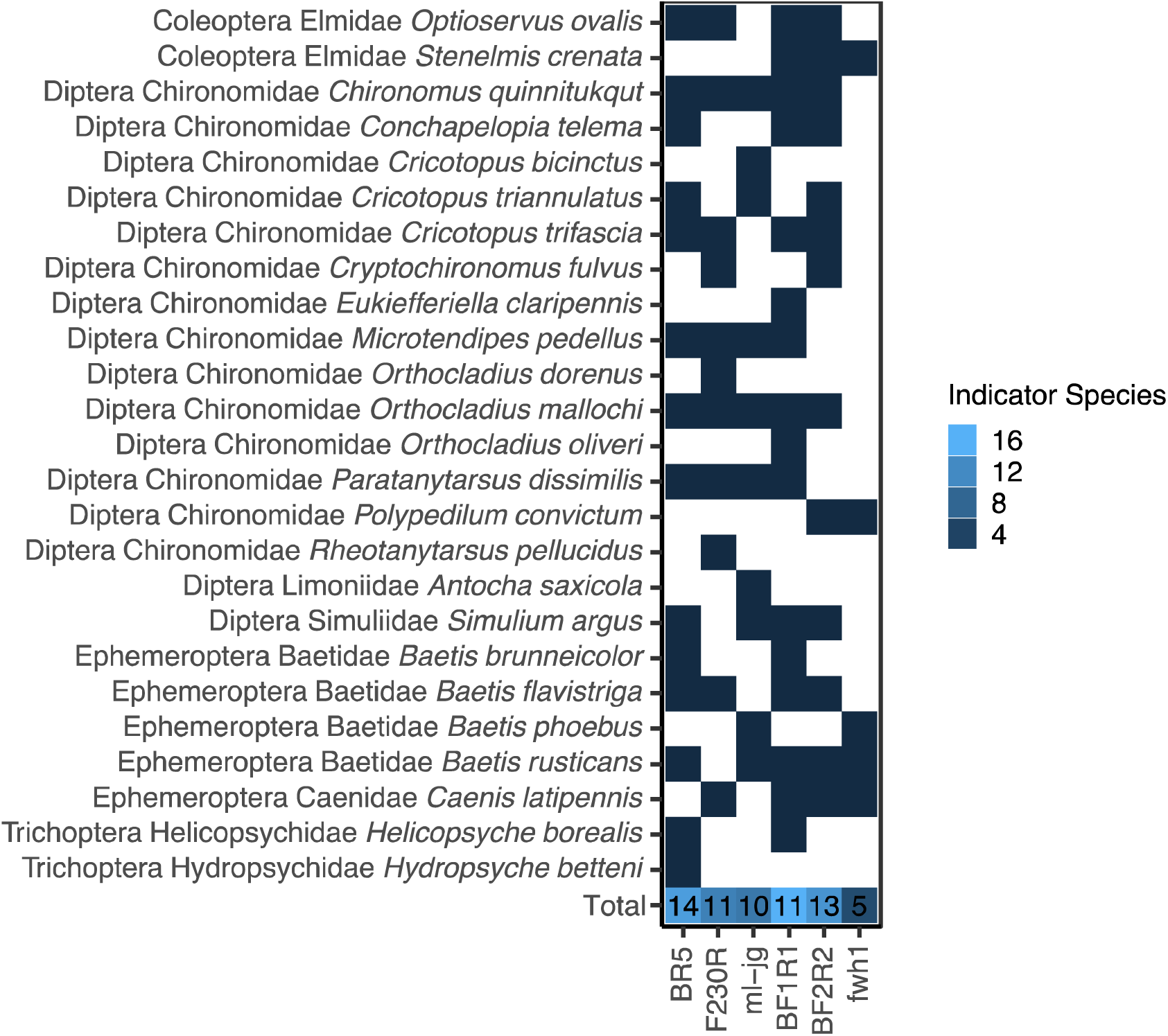
Site indicator taxa chosen based on metabarcode sequencing are comprised of Coleoptera, Diptera, Ephemeroptera, and Trichoptera. Presence is indicated by a dark square, absence by a white square. The total number of broadscale indicator taxa detected by each amplicon is shown in the bottom row according to the legend.

To investigate the effect of amplicon choice on beta diversity we looked at how sites cluster with respect to COI amplicon choice and PCR replicates. We compared all the data at the ESV rank (Figure 5). Samples cluster by site (stress = 0.154, linear R^2^ = 0.912). We found significant heterogeneity of beta diversity among sites (p-value < 0.05), but since we had a balanced design, proceeded to use PERMANOVA to test the significance of groupings (Anderson & Walsh, 2013). There were no significant interactions among sites, makers, or PCR replicates. Amplicon choice and PCR replicate did not explain any significant variation in beta diversity among samples, but sites explained 76% of the variation among samples (R^2^ = 0.76, p-value = 0.001).

**Figure 5.**
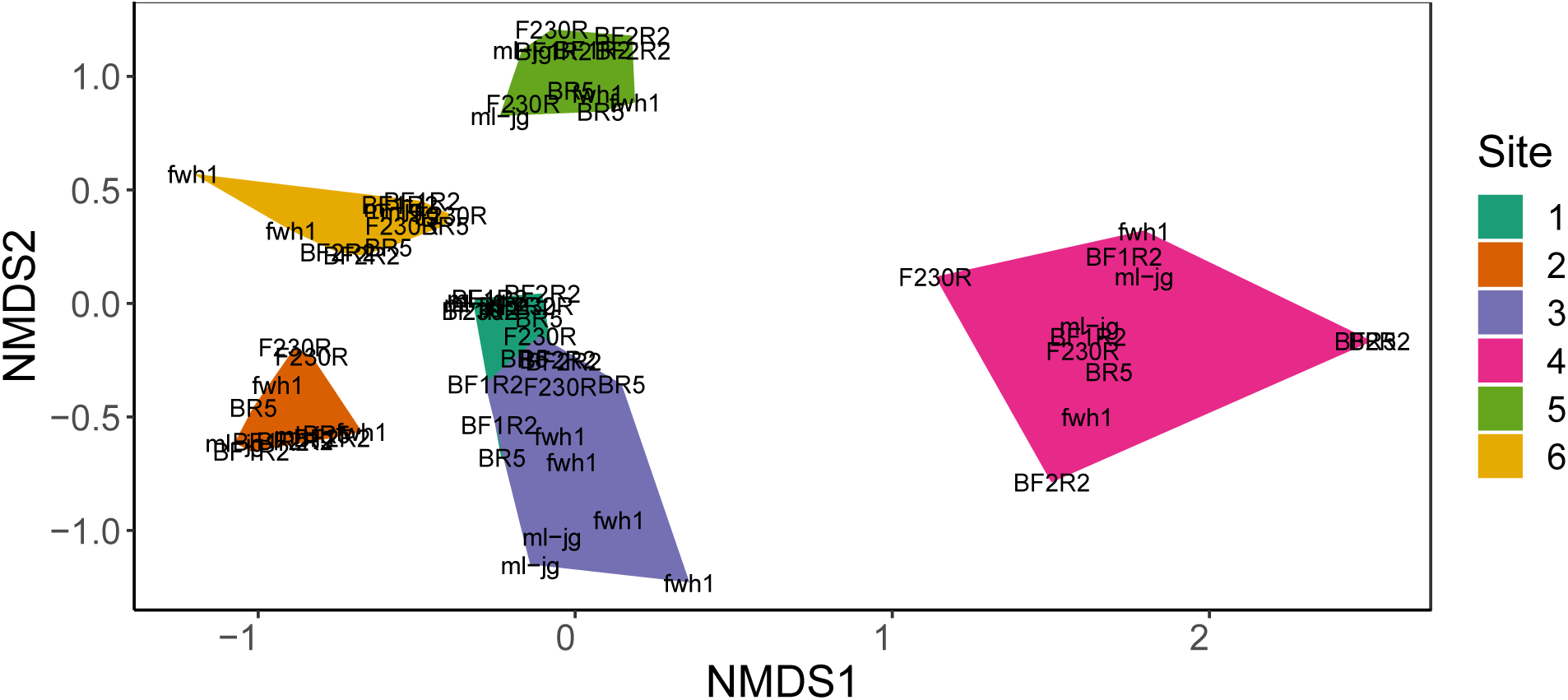
Samples cluster mainly by site despite differences in amplicons and replicates. Results are based normalized data. COI amplicons are labelled directly in the plot. Amplicons shown twice represent the two PCR replicates. Sites are grouped by color according to the legend.

## Discussion

As showcased in recent literature, DNA metabarcoding has gained significant popularity in various ecological studies where biodiversity in a habitat or a sample is investigated (Bik et al., 2012; Yu et al., 2012; J. Gibson et al., 2014, 2015; Creer et al., 2016; Leese et al., 2016; Bush et al., 2017; Porter & Hajibabaei, 2018c; Bush et al., Submitted). In this study we show that the optimal choice of amplicon(s) ought to be based on the objective of the study: optimizing richness, optimizing the differentiation of samples based on sites/conditions, or optimizing the detection of target taxa. Here we show the impact of using varied primer sets all of which have been used in recent metabarcoding studies of freshwater benthic macroinvertebrates.

As predicted, different primer sets produced varied richness results. For example, while the ml-jg amplicon produced the highest overall ESV richness, combinations of amplicons together detected even greater richness. Moreover, even though ml-jg maximizes ESV richness, at the species rank the best choice is F230R, yet at the genus rank BR5 optimizes richness. The decision to present the results of a study at various taxonomic ranks is often based on the desire to include all the data (ESV rank), or to present results at a fine level of taxonomic resolution (species rank), or to present results based on previous knowledge. For example, 94% of North American freshwater specimens identified by morphology are represented by a DNA sequence so it may be desirable to present results at the genus rank (Curry, Gibson, Shokralla, Hajibabaei, & Baird, 2018). These observations have important implications for choosing primers, especially when considering the level of standardization required in biomonitoring programs. While the use of a single primer set is desirable, richness based on metabarcoding is sensitive to the number of combined amplicons, primer choice, and the taxonomic resolution of the results. Based on results from this study and elsewhere, primer binding biases during amplification steps can have tangible impacts on results and using multiple primer sets will aid in increasing taxonomic coverage (J. Clarke et al., 2009; Bellemain et al., 2010; Hajibabaei, Spall, Shokralla, & van Konynenburg, 2012; J. Gibson et al., 2014). For the sake of flexibility and forward compatibility, aside from the storage of raw data, we encourage data reporting at the ESV rank. When reports are summarized to other taxonomic ranks, we encourage disclaimer statements that results are limited by the taxonomic coverage of current reference databases that may improve in the future (Porter & Hajibabaei, 2018b).

Our study provides important insights with regards to use of varied PCR primer sets and replicates. Contrary to measures of alpha diversity (above), beta diversity measures do not seem to be affected by primer sets or PCR replicates when ESVs are used for the spatial analysis. In other words, spatial separation of sites based on these varied parameters are robust as used in biomonitoring applications. While the use of standard primer sets may be desirable, it may not be required as beta diversity is less sensitive to primer choice and technical replicates.

Another important and widespread use of metabarcoding data is in determining ecosystem status or “biomonitoring” where the state of the ecosystem is derived from bioindicator assemblages such as EPTC (Buss et al., 2002). The recovery of bioindicators from metabarcoding data in this study varied with amplicon choice. For example, the BF1R2 amplicon detects the greatest number of broadscale Arthropoda site indicators but the F230R amplicon detects the greatest number of traditional EPTC site indicators. This has implications for benchmarking studies that compare metabarcoding against the results based on the use of traditional bioindicators. With the application of DNA-based methods, our ability to detect a broad range of taxa has improved such that it may not be necessary to limit bioindicator reporting to just the traditional bioindicators (Bush et al., Submitted).

Note that even though equimolar amounts of each amplicon were combined for sequencing, variable numbers of reads were obtained across amplicons. This may be caused by variable amplification efficiency during library preparation or slight differences in the number of transferred amplicons when they are pooled prior to library preparation. Since the recovery of variable library sizes is such a common occurrence, it is important to normalize library size across samples prior to conducting data analysis. It has been shown that there is a trade-off between the use of rarefaction (removal of sequences such that each sample can be compared at a common library size) to reduce the false positive rate, and a loss of sensitivity because of the removal of sequences (McMurdie & Holmes, 2014; S. J. Weiss et al., n.d.). This has implications for beta diversity analyses, where false positives can occur when samples cluster by sequencing depth obscuring real differences, especially for samples with very small library sizes.Common normalization methods include rarefaction down to the smallest library size and working with proportions (ESV reads per sample / total reads per sample). A simulation study showed that rarefaction combined with the analysis of presence-absence data worked best to cluster samples when groups are substantially different (S. Weiss et al., 2017). For differential abundance testing, however, methods that take into consideration the compositional nature of metabarcode datasets (log ratio transformation) may be more appropriate (Gloor, Macklaim, Pawlowsky-Glahn, & Egozcue, 2017; S. Weiss et al., 2017; S. J. Weiss et al., n.d.).

## Conclusions

This study analyzed how arthropod richness, beta diversity, and recovery of site indicator taxa vary with COI amplicon choice. We show how richness is sensitive to primer choice and the combined use of multiple COI amplicons; beta diversity is robust to primer choice and PCR replicates when ESVs are used for this analysis; and that limiting analyses to the traditional bioindicator assemblages may not capture the diversity of broadscale arthropod site indicators that can be recovered from COI metabarcoding. We note that the proportion of raw reads retained in the final set of Arthropod ESVs was relatively low (13%) and this proportion varied noticeably among different primer sets. There are several reasons why raw reads are filtered out of the final dataset: the removal of chimeras during PCR, removal of sequences with errors generated during PCR or sequencing, and/or to the removal of the substantial ‘tail’ of rare taxa as commonly seen in such studies reflecting either genuine rare novelty or artefacts (Kunin, Engelbrektson, Ochman, & Hugenholtz, 2010; Brown et al., 2015). Regardless, we suggest that method efficiency may also be gauged using a similar measure of raw data that can be retained in final analyses.

### Data Accessibility

Raw reads were submitted to the NCBI SRA: #. FASTA files of the final ESVs are available as supplementary material. The SCVUC semi-automated bioinformatic pipeline is available from GitHub (#). The custom scripts used to generate figures are also available from GitHub (#).

## Acknowledgements

This work was funded by the government of Canada through Genome Canada and the Ontario Genomics. TMP would like to acknowledge funding from the Canadian government through the Genomics Research and Development Initiative, inter-departmental Ecobiomics project.

