## Supporting Information for "COI metabarcoding primer choice affects richness and recovery of indicator taxa in freshwater systems"

**Table S1: Study sites**

| Sample | Region | Collected | Lat | Long |
|---|---|---|---|---|
| 1 | Laurel | May 31, 2018 | 43°28'2.64"N | 80°31'59.33"W |
| 2 | Clair | May 29, 2018 | 43°27'46.93"N | 80°32'56.75"W |
| 3 | Laurel | May 29, 2018 | 43°28'39.31"N | 80°33'30.45"W |
| 4 | Beaver | May 15 2018 | 43°29'28.24"N | 80°37'27.80"W |
| 5 | Laurel | May 17 2018 | 43°28'49.40"N | 80°36'15.37"W |
| 6 | Claire Lake Outflow | May 25, 2018 | 43°27'47.27"N | 80°33'7.83"W |

**Table S2: Reads counts for all taxa**

| | BR5 | F230R | ml-jg | BF1 | BF2 | fwh1 | Total |
|---|---|---|---|---|---|---|---|
| Raw | N/A | N/A | N/A | N/A | N/A | N/A | 9,980,584 x 2 |
| Paired | N/A | N/A | N/A | N/A | N/A | N/A | 8,253,974 |
| Primer trimmed | 1,113,306 | 1,523,753 | 1,492,705 | 1,774,254 | 941,889 | 773,201 | 7,619,108 |

N/A – Not applicable, as the amplicons were pooled before sequencing, then sorted by primer sequence at the primer trimming step

**Table S3: ESV counts for all taxa**

| | **BR5** | **F230R** | **ml-jg** | **BF1R2** | **BF2R2** | **fwh1** | **Total** |
|---|---|---|---|---|---|---|---|
| ESVs | 3,494 | 2,605 | 3,305 | 6,139 | 3,491 | 1,944 | 20,978 |
| Reads in ESVs | 324,721 | 643,405 | 375,119 | 432,848 | 80,886 | 457,968 | 2,314,947 |
| Proportion of raw reads (%) | 3.3 | 6.4 | 3.8 | 4.3 | 0.8 | 4.6 | 23.2* |

* ~ 77% of raw reads removed during denoising (putative sequence errors, chimeras, PhiX contamination, rare singletons and doubletons

**Table S4: Filtering for high confidence Arthropoda identifications affects the proportion of assignments retained across taxonomic ranks**

| | **ESVs** | **Species** | **Genus** | **Family** | **Order** | **Class** |
|---|---|---|---|---|---|---|
| High confidence assignments* | 4,940 | 120 | 100 | 69 | 44 | 9 |
| All unique assignments | 4,940 | 1049 | 660 | 270 | 44 | 9 |
| Proportion of assignments retained after applying bootstrap support cutoffs (%) | 100 | 11.4 | 15.2 | 25.6 | 100 | 100 |
| Reads in high confidence assignments | 1,280,397 | 718,120 | 775,884 | 1,012,057 | 1,280,397 | 1,280,397 |
| Proportion raw reads in high confidence assignments (%) | 12.8 | 7.2 | 7.8 | 10.1 | 12.8 | 12.8 |

* ESVs not taxonomically assigned so no bootstrap support filtering required; Species >= 0.70 bootstrap support cutoff; Genus >= 0.30; Family >=0.20; No bootstrap support filtering needed at the order or class ranks to ensure 99% correct assignments (95% for species)

**Figure S1. Sequencing depth is saturated for most samples.** For each primer, color coded as in the legend, there are 12 lines for the 6 field sampling sites x 2 PCR replicates. The vertical line indicates the 15th percentile library size that was used to normalize variable library sizes for subsequent diversity analyses. ESVs = exact sequence variants.

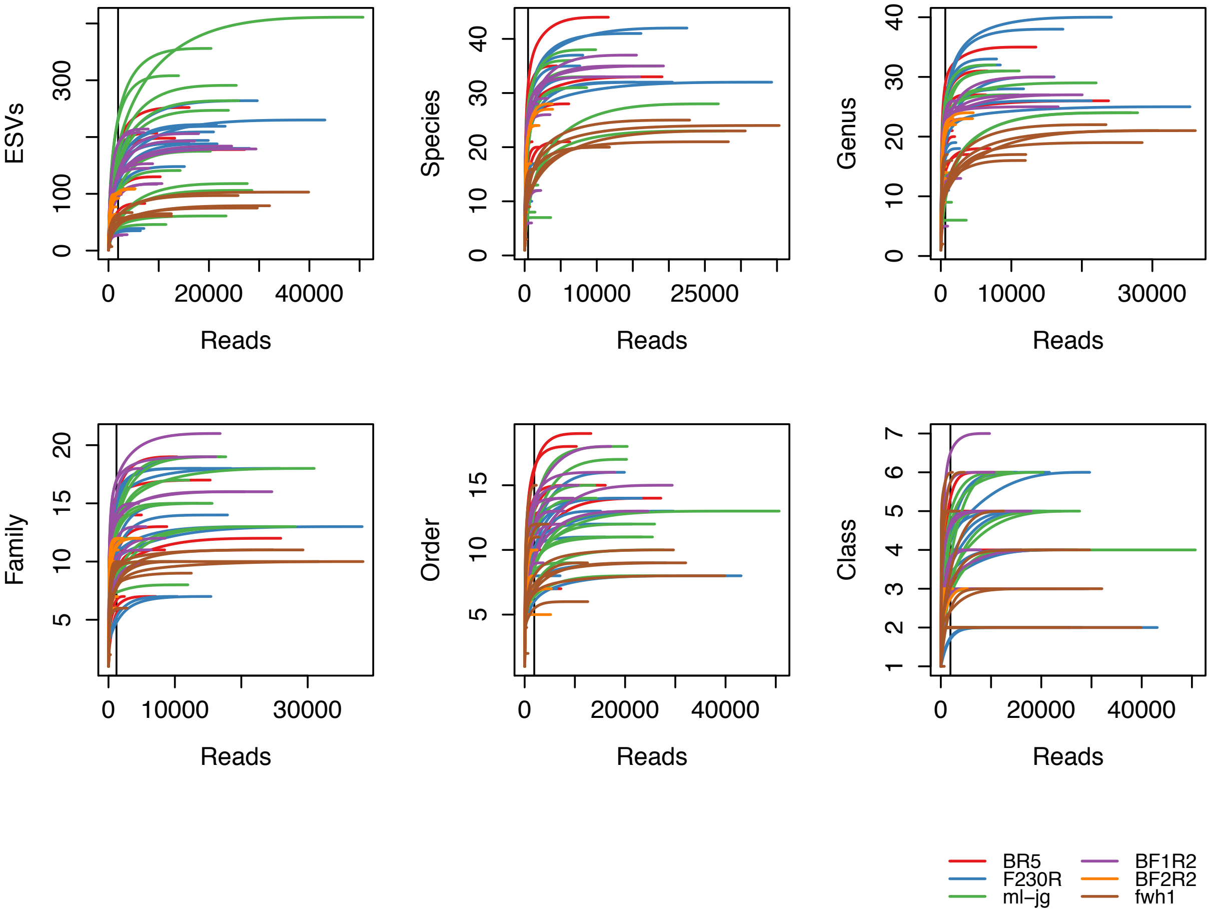

**Figure S2. Arthropoda taxa, especially the target assemblage comprised of Ephemeroptera, Plecoptera, Trichoptera, and Chironomidae, form a large proportion of the detected community.** Part A) shows the proportion of the top 10 phyla and the count value for the top 3 phyla are labelled in the plot. Part B) shows the proportion of all Arthropoda that are EPTC (Ephemeroptera, Plecoptera, Trichoptera, Chironomidae). Results based on raw data before normalization.

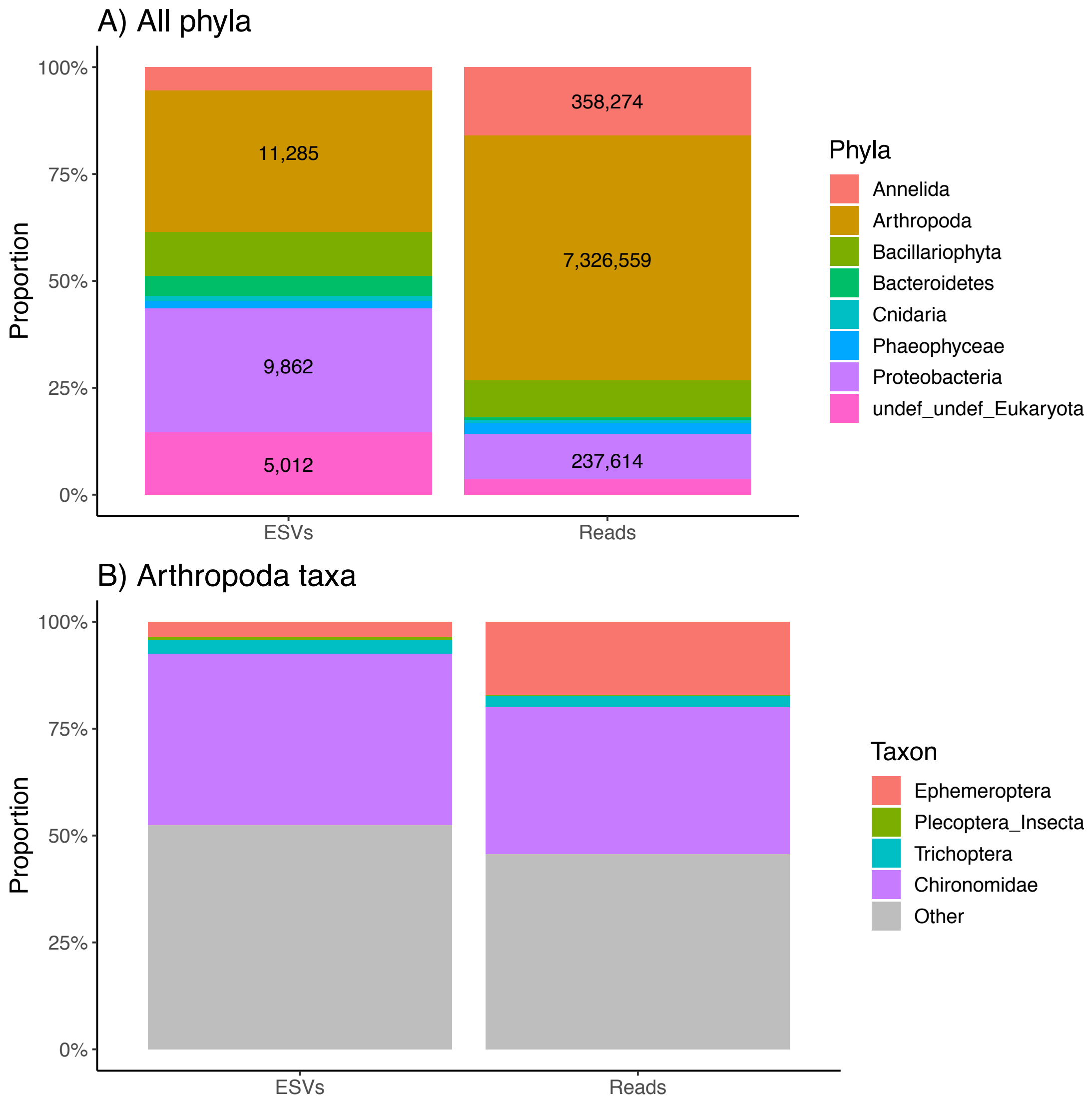

**Figure S3. Most primers recovered similar average Arthropoda richness across sites.** The first panel shows the 6 COI amplicons tested. Based on normalized data. Results shown are for 2 pooled PCR replicates at the ESV rank. ESV = exact sequence variant.

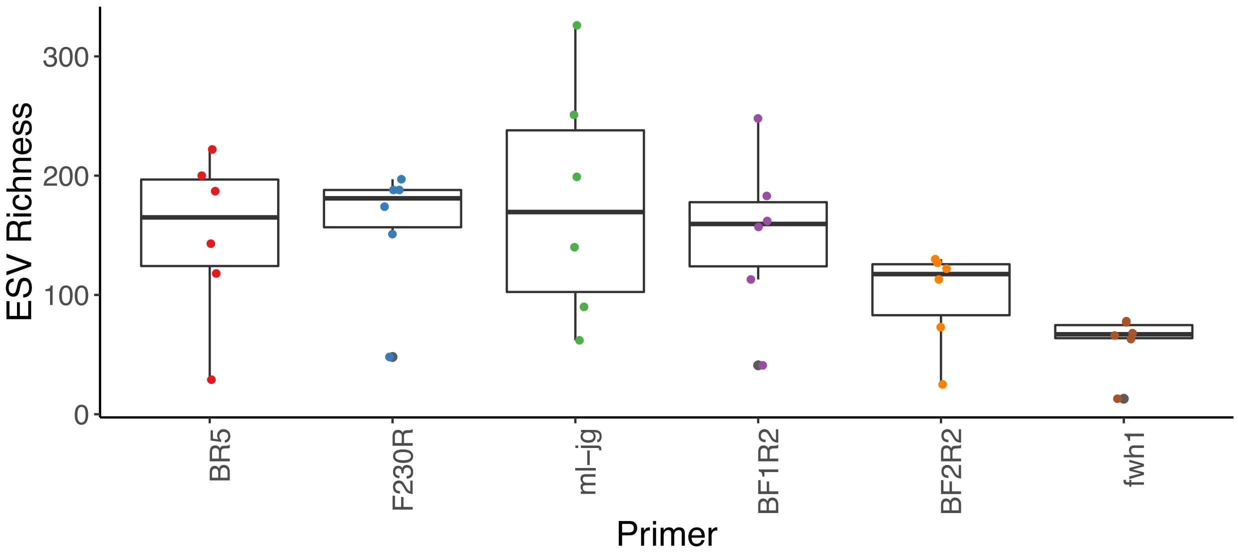

**Figure S4. The primer that detects the highest Arthropoda richness varies by rank.** Richness from each COI amplicon at a variety of taxonomic ranks are shown. Results are based on normalized data. ESV = exact sequence variant.

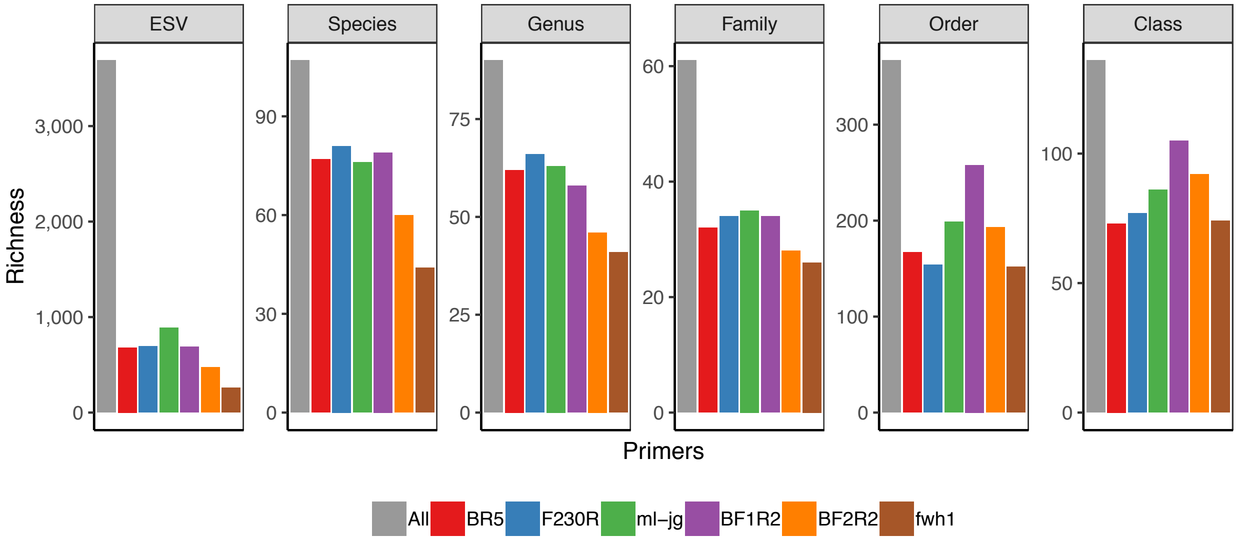

**Figure S5. Arthropoda ESV presence-absence is positively correlated between the soil and tissue DNA extraction kits.** Pearson correlation coefficients are shown. Results are based on normalized data. ESV = exact sequence variants. Label naming convention is as follows: DNA extraction kit _ amplicon _ site _ PCR replicate. A = BR5, B = F230R.

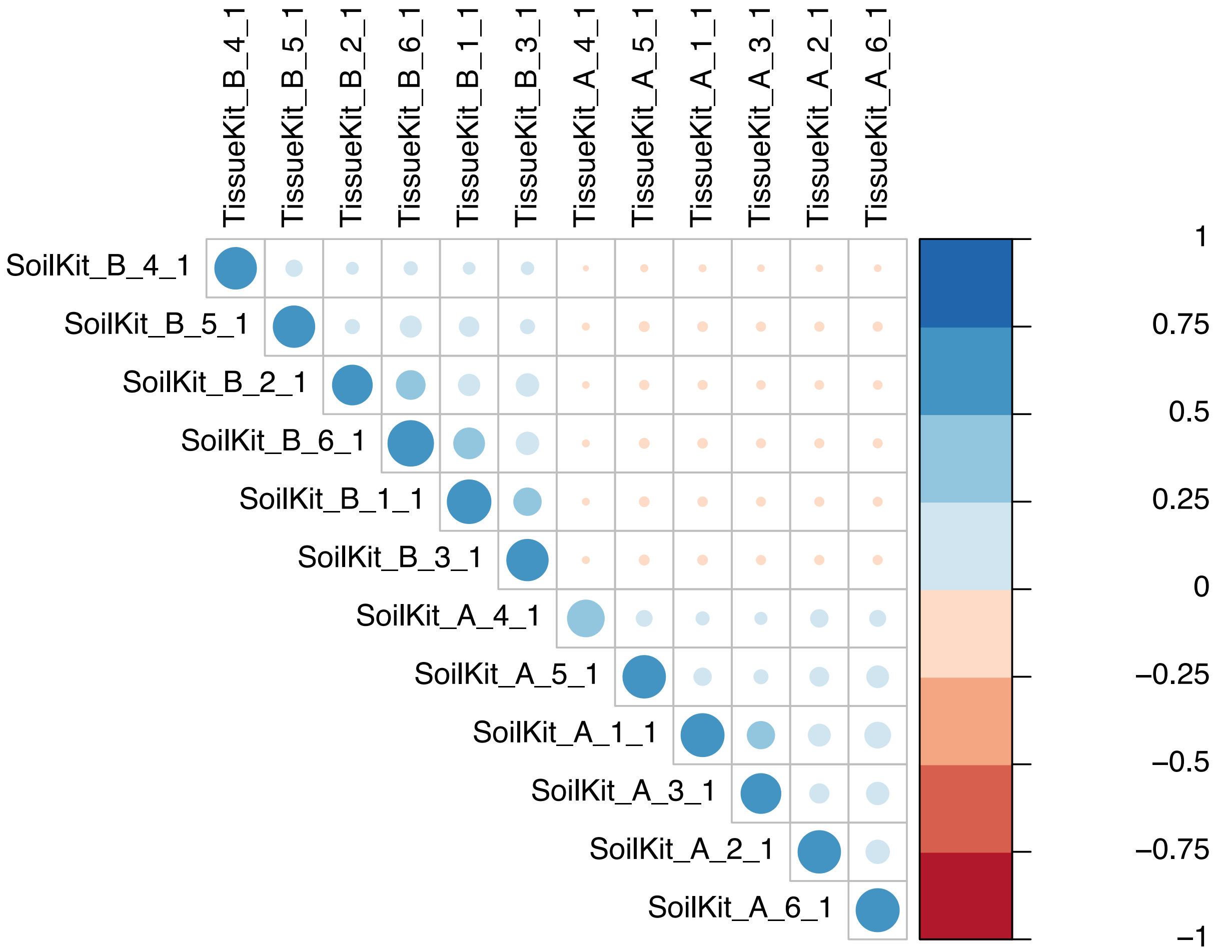

**Figure S6. Arthropoda ESV presence-absence is positively correlated between the first and second PCR replicate.** Pearson correlation coefficients are shown. Results are based on normalized data. ESV = exact sequence variants. Label naming convention is as follows: DNA extraction kit _ amplicon _ site _ PCR replicate. A = BR5, B = F230R, C = ml-jg, D = BF1, E = BF2, F = fwh1.

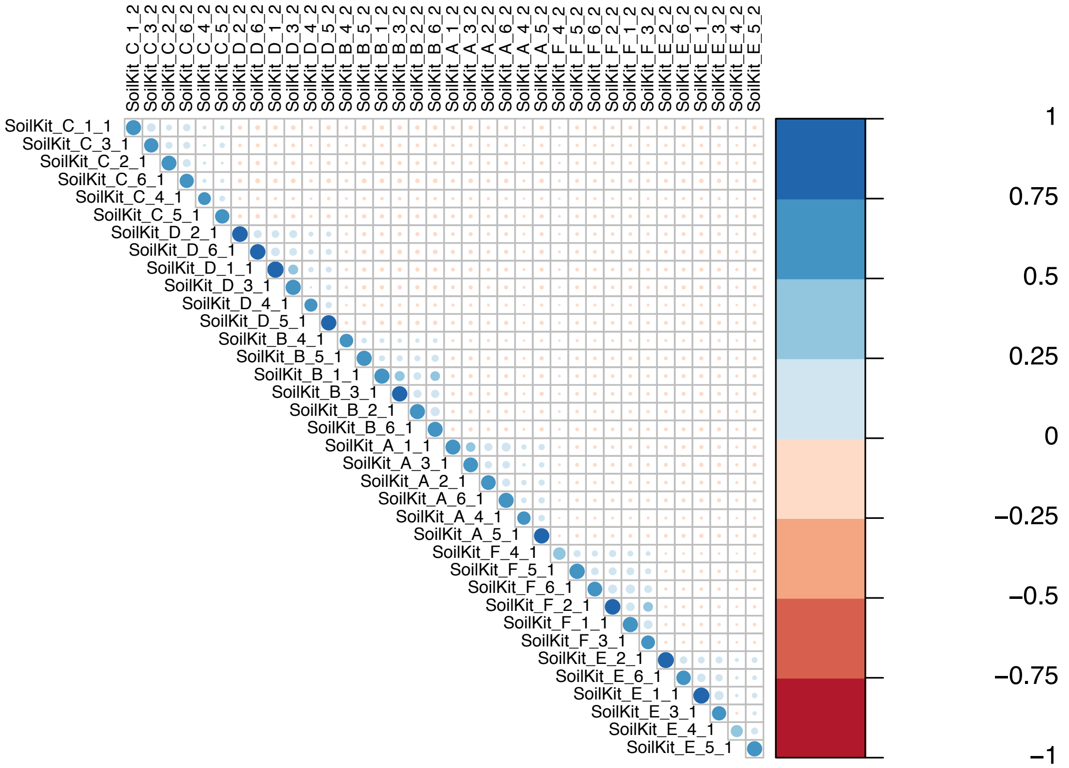
